# A Multi-Layered Computational Structural Genomics Approach Enhances Domain-Specific Interpretation of Kleefstra Syndrome Variants in EHMT1

**DOI:** 10.1101/2023.09.06.556558

**Authors:** Young-In Chi, Salomão D. Jorge, Davin R. Jensen, Brian C. Smith, Brian F. Volkman, Angela J. Mathison, Gwen Lomberk, Michael T. Zimmermann, Raul Urrutia

## Abstract

This study investigates the functional significance of assorted variants of uncertain significance (VUS) in euchromatic histone lysine methyltransferase 1 (EHMT1), which is critical for early development and normal physiology. EHMT1 mutations cause Kleefstra syndrome and are linked to various human cancers. However, accurate functional interpretation of these variants are yet to be made, limiting diagnoses and future research. To overcome this, we integrate conventional tools for variant calling with computational biophysics and biochemistry to conduct multi-layered mechanistic analyses of the SET catalytic domain of EHMT1, which is critical for this protein function. We use molecular mechanics and molecular dynamics (MD)-based metrics to analyze the SET domain structure and functional motions resulting from 97 Kleefstra syndrome missense variants within this domain. Our approach allows us to classify the variants in a mechanistic manner into SV (Structural Variant), DV (Dynamic Variant), SDV (Structural and Dynamic Variant), and VUS (Variant of Uncertain Significance). Our findings reveal that the damaging variants are mostly mapped around the active site, substrate binding site, and pre-SET regions. Overall, we report an improvement for this method over conventional tools for variant interpretation and simultaneously provide a molecular mechanism of variant dysfunction.

## INTRODUCTION

Over the last three decades, extensive work has recognized that histone modifications are central to epigenetic regulation. Epigenetic dysregulation caused by mutations in components of histone-modifying enzymes leads to various human diseases known as “Chromatinopathies” (1). EHMT1, also called G9a-like protein (GLP), catalyzes mono- and di-methylation of Lys9 of histone H3 (H3K9me1 and H3K9me2) for gene silencing (2). EHMT1 alterations are associated with Kleefstra syndrome (OMIM 610253), a neurodevelopmental disorder, and different tumor types, including uterine, adrenocortical and skin melanoma, and stomach adenocarcinoma (3). To benefit patients with cancer and suspected Chromatinopathies, improved methods are necessary to interpret EHMT1 genetic alterations.

Belongs to the SET domain-containing methyltransferase family, the human EHMT1 protein consists of 1298 amino acids, characterized by distinctive domains, including the transactivation domain at the *N*-terminus, the cysteine-rich domain, the ankyrin repeat domain with scaffolding function, and the enzymatic SET domain at the *C*-terminus. Responsible for the writer function of EHMT1, the SET domain catalyzes mono- and di-methylation of the H3K9 residue. Reader function is conferred by the ankyrin repeat domain, which recognizes the same histone modification (H3K9me1/2) and mediates higher-order complex formations for gene and epigenome regulations (4). Thus, although the full-length protein cohesively exerts its function, the impact of mutations on domain-specific molecular mechanisms should be considered, as each domain represents an independent folding unit and possesses a discrete function. While numerous Kleefstra syndrome germline variants have been identified from patients (5, 6), their pathogenicity and dysfunctional molecular defects are poorly understood. Current variant interpretations rely heavily on 2D sequence-based information and limited structural and functional data (7). Thus, computational approaches and multiplexed experimental data are urgently needed to improve prediction power and delineate potential dysfunctional mechanisms (8).

This study reports a domain-wide analysis of SET catalytic domain variants associated with the Kleefstra syndrome congenital disease to fill this knowledge gap. The SET catalytic domain variants have unique molecular features that can be shared by its homolog EHMT2 and other members of the SET domain-containing methyltransferases. We studied 97 missense variants (on 82 residues) within the EHMT1 SET domain using the available crystal structure (PDB access code 3HNA) to predict their mutational impacts on the structural and dynamic properties of the protein. We applied selective analytical tools from each protein layer representing universal or protein-specific and global or local considerations, including folding/stability energy, structure perturbation, binding energy calculations, local geometry analyses, and all-atom MD simulations. These multi-tiered mechanistic-based analyses complement existing prediction tools and further enhance the mutational impact assessments highly relevant to protein structure and function. Thus, this study represents a novel approach to understanding the functional effects of these alterations by providing a broader characterization of genomic variants with dynamic modeling specific to a rare disease-associated SET domain.

## RESULTS

### Defining the mutational germline landscape associated with the causality of Kleefstra syndrome

Kleefstra syndrome is a rare genetic disorder caused by mutations in EHMT1. Many of these alterations are de novo mutations (5, 6). To study the impact of SET domain missense mutations, we extracted all EHMT1 missense variants found in patients diagnosed with Kleefstra syndrome from ClinVar, the most comprehensive public archive of genomic variations and interpretations of their relationships to diseases (9). Figure 1A shows the distribution, frequency, and database sources for all Kleefstra syndrome variants under study. We find no apparent ‘hot-spot’ regions, as variants are scattered across the entire sequence and 3D structure. Slightly higher alteration frequencies are found within the pre-SET and the core-SET domain, but the differences are insignificant. Figure 1B shows the molecular structure of the EHMT1 SET domain, and Figure 1C shows the variant mapping onto the molecular structure and their current pathogenicity annotations in ClinVar. Although professionals routinely use these database annotations as variant interpretation guidelines, most Kleefstra syndrome variants are still classified as variants of unknown significance due to insufficient evidence and incomplete impact assessment.

**Figure 1.**
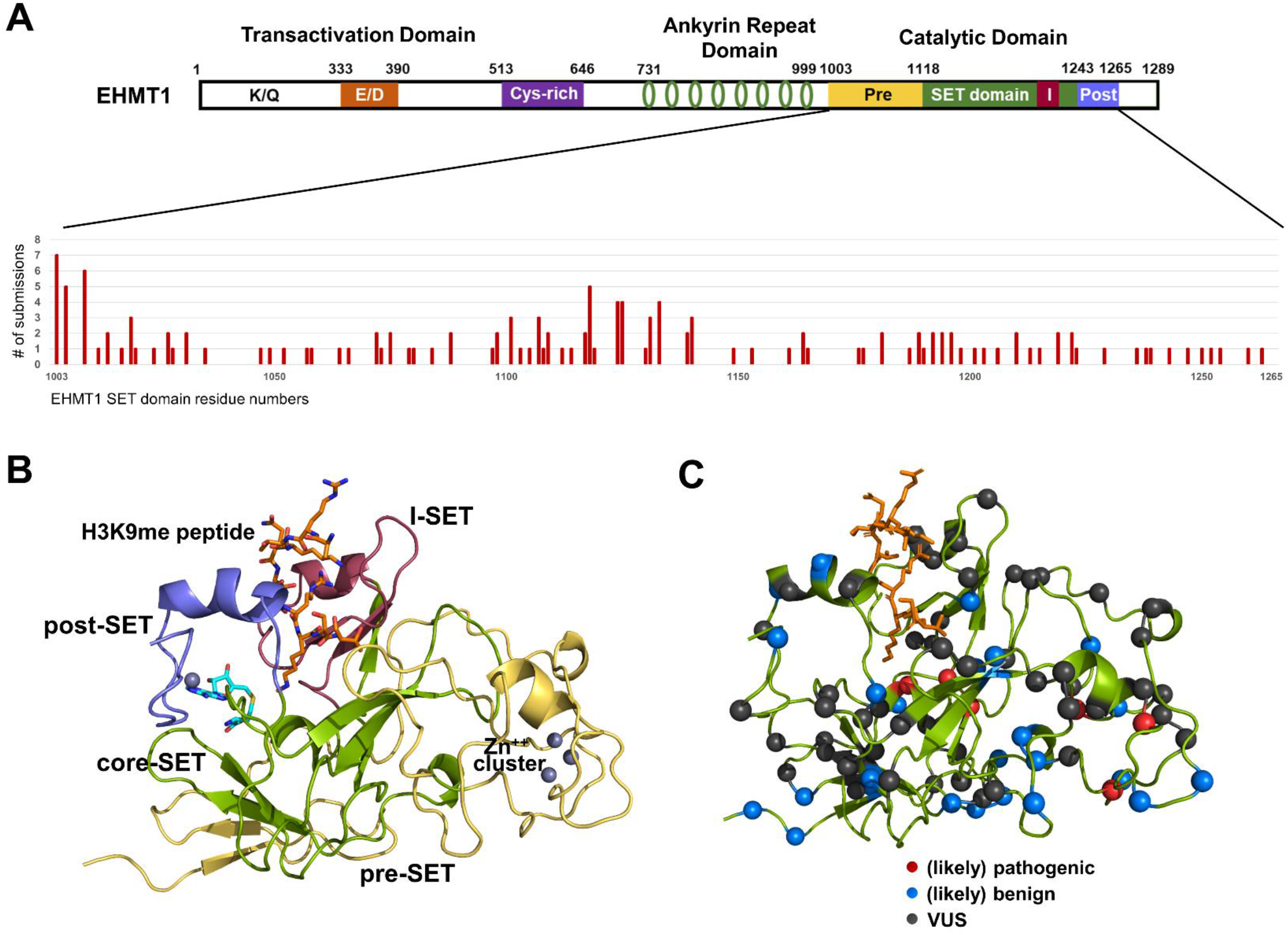
Protein architecture and Kleefstra syndrome variants of EHMT1. (A) Domain structure of EHMT1 and the distribution of Kleefstra syndrome missense variants within the SET domain. The cases are tabulated by the number of independent missense mutations found on a particular residue and reported to ClinVar. (B) Molecular structure of the EHMT1 SET domain. The same color codes for the sub-domains shown in (A) are used. The bound H3K9me peptide and the SAH cofactor are depicted as a ball-and-stick model. The structural zinc ion cluster is also indicated. (C) Current annotations of the variants under study in ClinVar and mapping of the 97 Kleefstra syndrome variants onto its molecular structure. Over 61% of the variants (60 out of 97), 32.0% (31 out of 97), and 6.2% (6 out of 97) are currently classified as VUS, (likely) benign, and (likely) pathogenic, respectively. Among these, 30 variants have also been identified as cancer somatic variants. These mutations are distributed over the entire sequence and its molecular structure, with slightly higher frequencies in the pre-SET and the core SET domains.

Among the 97 SET domain variants, 6 variants are currently annotated as pathogenic or likely pathogenic, and 31 are annotated as benign or likely benign in ClinVar (Fig. 1C). Among the pathogenic variants, C1073Y and R1197W have been biochemically characterized and proven to be damaging (10) and thus serve as positive controls for this study. In addition, among the benign variants, S1004N and V1006M have relatively high allele frequencies (> 1.0 × 10^−4^) in the healthy population gnomAD database (11) and are used as potentially tolerated or neutral variants for this study. These S1004N and V1006M variants are also indicated as SNP in the Single Nucleotide Polymorphism Database (dbSNP) and are expected to have no appreciable deleterious and pathogenic effects. Among the 97 variants, 30 have been observed somatically in human cancers (12). While EHMT1 and EHMT2 are commonly altered in human cancers, Kleefstra syndrome variants are only found in EHMT1. Thus, this set of variants represents EHMT1-specific unique germline variants and more common disease variants.

### Determining the functional dynamic motions that characterize the molecular fitness of the EHMT1 SET domain

The EHMT1 SET domain is divided into sub-domains – the canonical core-SET, pre-SET, post-SET, and a small insertion within the core-SET domain architecture, termed I-SET (Fig. 1B). The post-SET and I-SET make the substrate recognition site. In contrast, the active site is primarily located within the core-SET domain. The pre-SET region contains the structural zinc ion cluster and a dimerization interface with either EHMT1 or EHMT2 for biological homo- and hetero-dimer functional units. The MD trajectories of the wild type in complex with the SAM cofactor and the structural zinc ions show a coordinated movement with high mobility in the pre-SET region, which provides a dimerization interface. At the same time, relatively rigid motions in the active and substrate binding sites are observed (Supplementary Fig. S1 and Supplementary Movie M1). Thus, the histone H3 tail binds to the pre-formed stable recognition site and readily presents its H3K9me1 substrate moiety into the active site.

### Comprehensive assessment of EHMT1 variants using conventional genomic tools and computational biophysics and biochemistry

We aimed to combine highly correlated impacting scores from each protein layer, including sequence, structure, and dynamics. Currently, annotations of genomic variants are primarily based on 2D sequence conservation/residue coevolution, the physicochemical property of the substituted amino acid, and local structure considerations such as secondary structures (7, 13). Commonly used 2D sequence-based variant calling methods include SNP&Go (14), PROVEAN (15), PolyPhen2 (16), Rhapsody (17), CADD (18), and REVEL (19). Combined annotators, such as CADD and REVEL, show better performance (20). However, we recognize that protein function is not solely determined by its chemical composition but also by its molecular structure’s spatial arrangement and dynamic nature. Therefore, we applied selective analytical measures that operate on both structural and dynamics to investigate the disruptive effect of each mutation. Specifically, we conducted a series of static or dynamic structure-based analyses using either the original crystal structure (stressed) or energy-minimized structure (relaxed). Initially, we applied metrics that are universal to proteins, such as folding energy, protein stability, global/local structural perturbation, energetic frustration, and dynamics-based analyses, including root mean square deviation (RMSD), root mean square fluctuation (RMSF), a radius of gyration (Rg), and solvent accessible surface area (SASA). Subsequently, we calculated correlations among these scores to identify more functionally relevant, thus evolutionally conserved metrics for overall damaging assessment (Fig. 2). We hypothesize that integrating 2D sequence-based scores with scores from protein 3D structure and 4D time-dependent dynamic behaviors for molecular fitness will enhance the prediction power of variant interpretation (21-24). Our study demonstrates the importance of considering protein-specific and mechanistic-based interpretations of variants for clinical recommendations.

**Figure 2.**
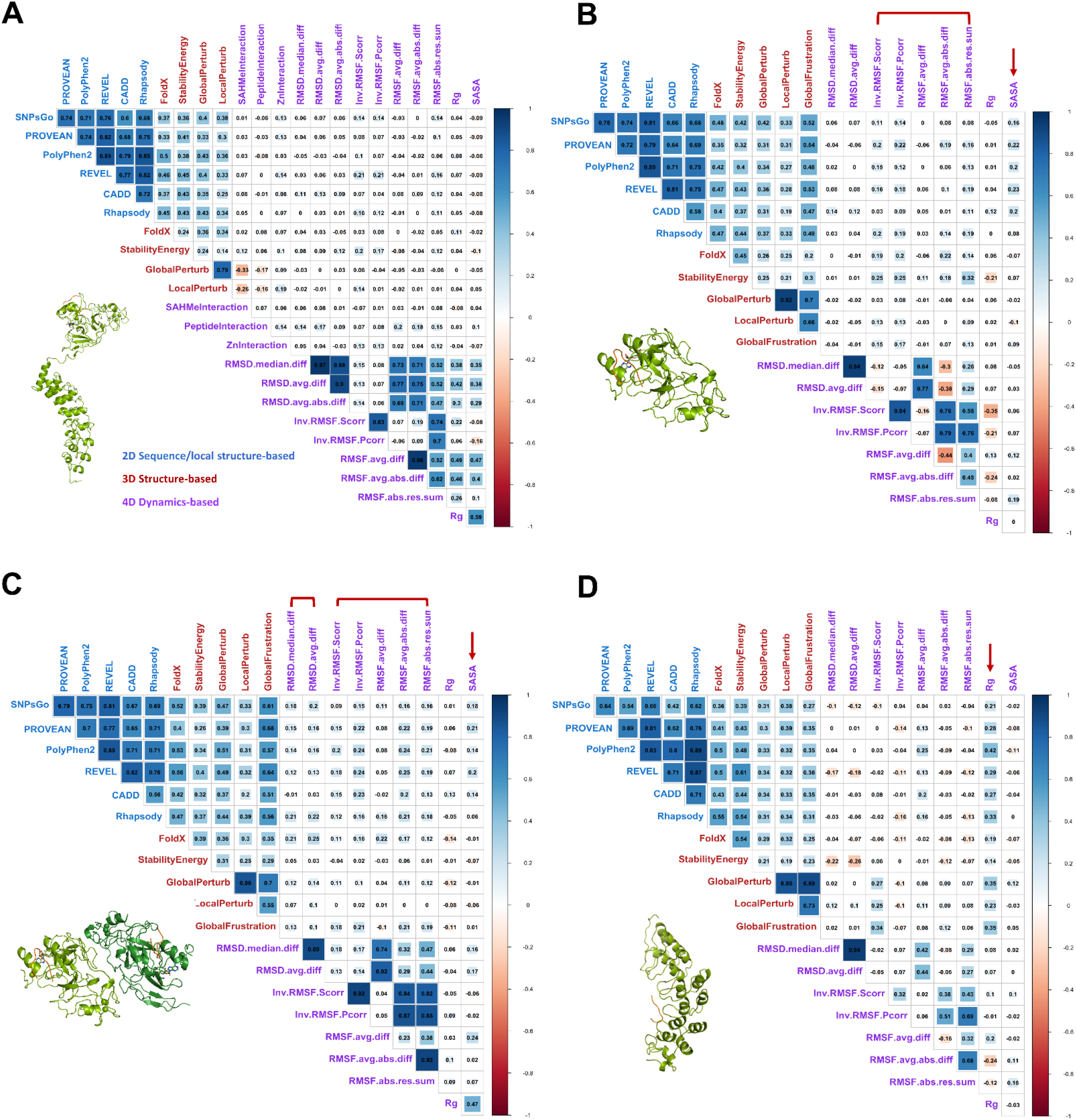
Identification of congruent and domain-specific MD-based metrics by the cross-correlation matrix of the individual damaging scores. (A) Cross-correlation matrix with various measures of universal and protein-specific metrics using the two-domain structure. Although all 3D structure-based scores show notable congruence with the 2D sequence-based scores, nothing emerged from the 4D MD-based scores. (B) Cross-correlation matrix with various measures of universal metrics using the SET domain-only structure. Among the 4D MD-based scores, RMSF-related scores (indicated by a bracket) and SASA (indicated by an arrow) show congruence with the sequence-based scores. (C) Cross-correlation matrix with various measures of universal metrics using the biological EHMT1:EHMT2 SET domain heterodimer structure. Among the 4D MD-based scores, in addition to RMSF-related scores and SASA, RMSD-related scores also show congruence, implying these features have been evolutionally conserved. (D) Cross-correlation matrix with various measures of universal metrics using the ankyrin repeat domain-only structure. Rg is a congruent and functionally relevant metric among the 4D MD-based scores.

### Analysis with the two-domain structure, congruence among individual scores, and unraveling the need to perform domain-wide analysis

We used various analytical measures that operate on both structural and dynamics to probe the disruptive effect of each mutation. We initially constructed a two-domain structure containing both SET and ankyrin repeat domains by superposing the overlapping region of individual domain structures (PDB access codes 3HNA and 6BY9). We then performed various calculations using universal metrics from each protein layer (Fig. 2A). We used the difference values between the wild type and the variants as potential damaging scores (24). We expected a correlation among the impact scores from all three layers of proteins, as protein sequence, structure, and dynamics are highly coupled, and their coupling strongly influences the evolutionary selection unique to each protein’s molecular function (25-27). The cross-correlation matrix calculated with the individual scores revealed that all structure-based scores showed noticeable congruence with sequence-based scores, reaffirming the interrelationship between protein structure and sequence and the effectiveness of structure-based metrics as universal metrics for all proteins (Fig. 2A). However, MD-based scores showed little congruence with the sequence-based scores, possibly due to differential sets of functionally relevant metrics for each domain with a unique function. We conducted a domain-wide impact analysis using individual domain structures to test this possibility.

### Developing domain-specific effective and functionally relevant metrics increases the yield of discovering damaging variants

Unlike the two-domain structure analysis, when we performed the individual domain-wide analyses, positive correlations showed up for MD-based scores although the actual correlation values were lower than the structure-based scores. All structure-based scores showed significant congruence with sequence-based scores as expected, and some dynamic-based scores showed notable agreement with other scores in a domain-specific manner. For the monomeric SET domain, RMSF-related scores and SASA showed congruence with the sequence-based scores (Fig. 2B). Despite weak correlations, these differences are quite striking, and we hypothesize they have implications. Although the functional relevance of SASA cannot be readily explained (perhaps, it is related to dimerization or some unknown protein-protein interactions), correlations with the RMSF-related scores are reminiscent of those of the KDM6A catalytic domain in our prior studies (24). Although this needs to be tested against many different enzymes, concerted fluctuating frequency of dynamic motions throughout the molecule might be a common property essential for all reactions to be optimally catalyzed and has been conserved, thus can be an effective measure of functional disruption.

When the biological heterodimeric SET domain was used as a starting model (28), in addition to RMSF-related scores and SASA, RMSD-related scores also showed noticeable congruence (Fig. 2C). This might be related to the relative orientation between the monomers which can play a critical functional role (Supplementary Fig. S2). However, dimerization interaction energies show weaker correlations with the sequence-based scores (Supplementary Fig. S3). On the other hand, for the alpha-solenoid ankyrin repeat scaffolding domain, only Rg was a congruent and effective metric for functional disruption (Fig. 2D). For EHMT1, each domain has a different set of functionally relevant and more effective metrics for functional disruption. When scored collectively, the true signals can be canceled out, producing low correlations with other scores (Fig. 2A). As a result, the MD-based damaging scores from the two-domain model can become less reliable. Thus, we used the individual domain structures to complete domain-specific and comprehensive molecular fitness analyses. In this article, we present the data with the catalytic SET domain; the results with the ankyrin repeat domain will be published later with validation data. The entire molecular fitness scores for the SET domain are provided in Supplementary Table S1, and their plots against the variants for each metric are shown in Supplementary Figure S4.

### Parametrizing SET domain-specific metrics extend the predictive power of evaluating damaging variant in enzymatic domains

The SET domain is well-known for its catalytic mechanism involving critical functional roles of the SAM cofactor and two tyrosine residues (29, 30). Tyr1155 aligns the substrate lysine for the methyl transfer reaction, while Tyr1242 enhances the electrophilicity of the departing methyl group of the SAM cofactor (Fig. 3A). Optimal enzymatic activity relies on local geometry and mutual interactions, which can be affected by disease-causing mutations. Thus, we measured time-dependent interaction energies to assess potential damaging impacts and monitored the distances between critical functional elements from the molecular dynamics (MD) trajectories. We also calculated the differences between the wild type and the variants for the peptide interaction energy (substrate binding). We observed congruence between the sequence-based scores and all essential interaction- and distance-related scores, except for the SAM cofactor and the substrate target Lys interactions, when we calculated cross-correlations between these values and other scores (Fig. 3B). Thus, in the overall impact scoring, we included these domain-specific metrics, in addition to the congruent universal metrics such as RMSF and SASA (Fig. 3C). This confirmed the functional relevance of these metrics and supported their inclusion in the overall impact scoring.

**Figure 3.**
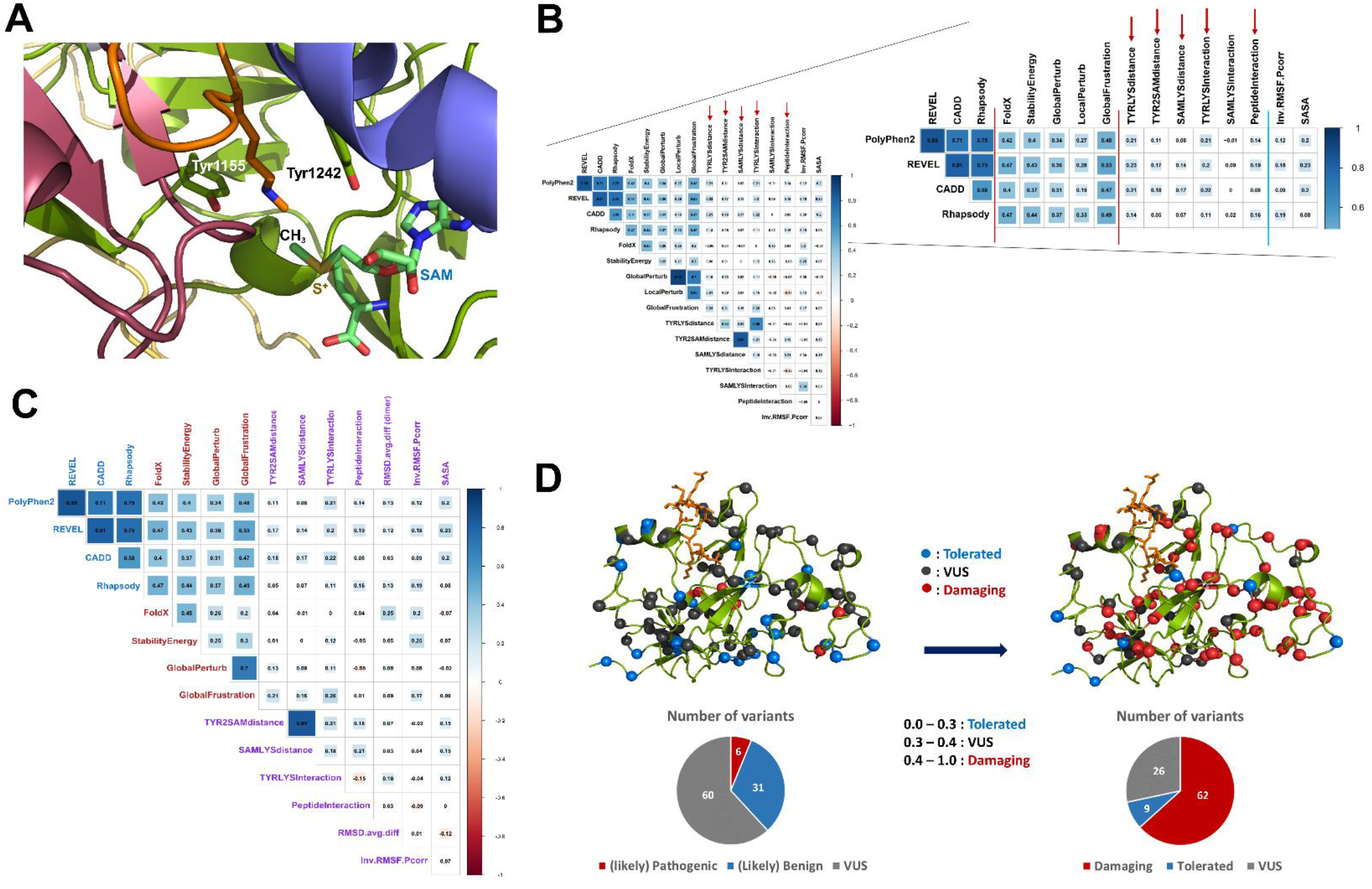
Identification of additional domain function-specific MD-based metrics, overall impact scoring, and reclassification of the variants. (A). Key functional elements in the active site. In addition to the SAM cofactor, two tyrosine residues play critical roles in catalysis. (B) Cross-correlation map with additional catalysis-related metrics and the universal MD-based metrics previously identified such as RMSF and SASA (separated by the cyan bar among the MD-based metrics). Noticeably congruent and functionally relevant domain-specific metrics are indicated by red arrows. Tyr and Tyr2 refer to the aligning Tyr1155 and the catalytic Tyr1242 residues, respectively. (C) Cross-correlation matrix of the scores from the finally chosen metrics for meta-score calculations that are concordant and functionally relevant, thus have been evolutionally conserved. (D) Mapping of pre- and post-classified EHMT1 Kleefstra syndrome variants. Meta-scoring reveals that the damaging variants (red) are concentrated near the substrate binding site, active site, and pre-SET region which contains the structural zinc-ion cluster and the dimerization interface, while tolerated variants (blue) are primarily found in the periphery or surface of the molecule. Pie charts at the bottom show the numbers of the pre- and post-classified variants in each category.

### Data integration and modeling allow meta-scoring and re-classification of EHMT1 SET domain Kleefstra syndrome variants

We selected four structure-based and seven dynamics-based metrics that show congruence with sequence-based scores and combined them with sequence-based scores to extend the predictive power of evaluating damaging variants in enzymatic domains (Fig. 3C). These metrics were used to compute an overall score for each variant, which can be numerically represented for practical use by clinicians and geneticists. However, calculating the final scores simply by summing up the individual scores is inappropriate, as the measurements for individual metrics are given in different units and have distinct ranges. Therefore, we chose to use Z-score scaling to transform the individual scores onto a zero to one range, commonly used by many sequence-based tools (31). We then averaged them for the final scores (Supplementary Table S1).

We used suggested thresholds for each prediction tool as guidelines to classify the variants. We re-classified the variants into three groups: tolerated (0-0.3), uncertain (0.3-0.4), and damaging (0.4-1.0) based on their overall damaging scores. The re-classified variants are shown in Figure 3D. This results in a balanced number of variants in each category. Out of 97 variants evaluated, 62 (63.9%) were classified as damaging, 9 (9.3%) were classified as tolerated, and 26 (26.8%) remained as variants of uncertain significance (VUS). However, this significantly improved from the 60/97 (61.9%) VUS identified in the current ClinVar annotations. The damaging variants are mostly found near the functional regions, while the tolerated variants are located in the periphery or molecular surface. Our analysis showed that the annotations of currently classified VUS have been improved using our method and our comprehensive structural genomics approach enhanced the prediction power of genomic variant interpretation.

### A molecular biophysics classification of variants and EHMT1/2 paralog analysis extends information on damaging effects on EHMT1 and generalizes results to related proteins

We further classified the damaging variants into structural (SV), dynamic (DV), and structural & dynamic variants (SDV) to provide mechanistic interpretations (24). We calculated molecular fitness scores by considering only structure- and dynamics-based scores, as shown in Supplementary Table S1. These molecular fitness evaluations revealed that 13 variants are expected to disrupt at least one of the structural features. In comparison, 51 variants are expected to disrupt at least one of the dynamic features (Fig. 4A). Among these, 11 variants disrupted both the protein’s structural and dynamic properties. The C1073Y and R1197W variants, previously characterized and proven to be damaging to protein function, are predicted to be damaging by our analysis (Fig. 4A left panel). In contrast, our analysis predicts two other variants from gnomAD with a relatively high allele frequency > 1.0 × 10^−4^ in general populations (S1004N and V1006M variants) to be tolerated by our analysis (Fig. 4A right panel). All previously annotated damaging variants are expected to be damaging by our analysis (Fig. 4B left pie chart). However, among the previous benign annotators, only 4 out of 31 are expected to be tolerated by our analysis, and 18 are expected to be damaging while 9 variants now belong to the VUS group (right pie chart). Thus, our results suggest that current annotations in the database tend to underestimate the damaging impact of genomic variants. The annotations of the currently classified VUS variants have also been improved using our method, and 38 out of 60 are expected to be damaging while 5 are expected to be tolerated (middle pie chart). Our analyses provide evidence that integrating 2D sequence-based scores with the scores from the protein 3D structure and 4D time-dependent dynamic behaviors for molecular fitness can enhance the prediction power of variant interpretation and provide potential molecular mechanisms for functional disruption.

**Figure 4.**
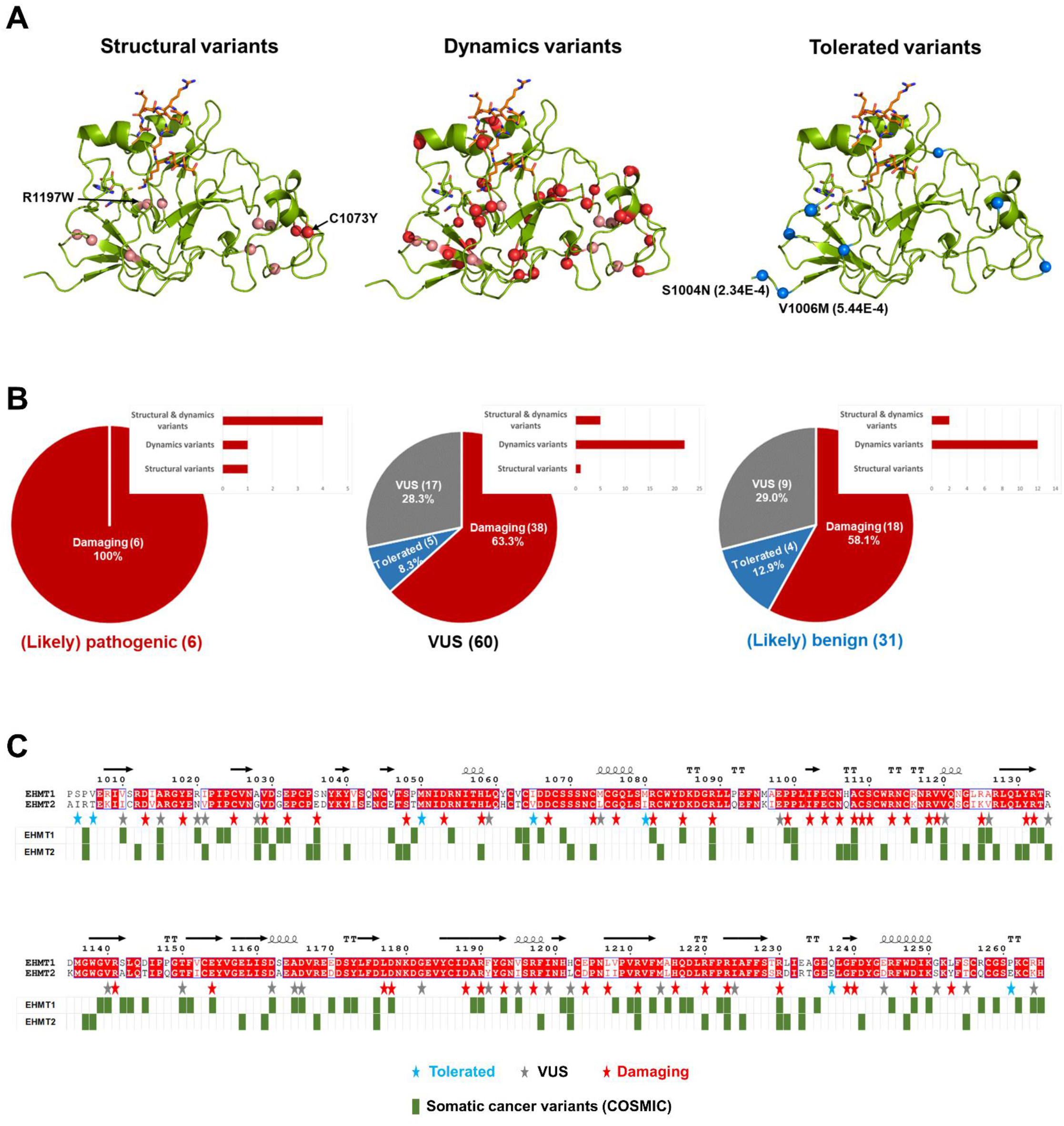
Further classification of the variants, comparison with pre-classification, and the EHMT1/2 paralog analysis. (A) Additional classification of the variants into structural variants (SV), dynamic variants (DV), and structural & dynamic variants (SDV), and mapping onto the molecular structure. SV or DV are indicated as red balls, SDVs are indicated in pink, and the tolerated varians are shown in blue. Previously characterized damaging variants (C1073Y and R1197W) and higher allele frequency variants among the healthy populations (S1004N and V1006M) are also indicated. (B) Comparison of pre-classification (ClinVar annotations) and post-classification by our molecular fitness analysis for each pre-classified group. We compare the two classification results using a pie chart that indicates damaging versus tolerated for our new classification results, for each of the three pre-classification categories. The inset bar chart shows the balance among the three damaging sub-categories. These mechanism-based interpretations should resolve the conflicting variants (middle) and provide enhanced interpretations for all variants. The numbers of the variants in each group are indicated in parentheses. (C) Comparison of post-classification and EHMT paralog annotation analysis. Sequence alignment of human EHMT1 and EHMT2 with indications of conserved residues (red boxes) and revised annotations (stars below). More than one variant is found on some of these residues. Cancer somatic variants in both proteins are also indicated at the bottom (green boxes). Residue numbers are based on EHMT1. Secondary structure elements are shown at the top of each sequence alignment. Positions of the re-classified variants of the current study are indicated by red (damaging), VUS (black), and blue (tolerated) stars at the bottom of aligned sequences. All germline variants on the conserved residues are predicted to be damaging by our comprehensive analysis, and none of the tolerated germline variants are represented in the cancer somatic variants.

Finally, we evaluated the results against the sequence conservation between EHMT1 and EHMT2. Our data indicated that most damaging variants (57/62, 91.9%) are found on canonical residues that are highly conserved for structural and functional reasons while most tolerated ones (8/9, 88.9%) are found in varying residues (Fig. 4C). None of the tolerated germline variants are represented in the cancer somatic variants. Cancer somatic variants are found throughout the sequence including many varying residues, some of which might represent polymorphisms and be tolerated. Overall, our data showed that the damaging consequences of these variants are well-represented in the human disease genomic landscape. This sequence information and mechanistic-based structural bioinformatics have the potential to provide better diagnosis, risk assessment, and clinical guidelines for observed variants within individualized medicine.

## DISCUSSION

Proteins perform vital functions by being made up of amino acids that fold into a unique 3D structure. The effect of genetic variation on protein structure and function can be dramatic, with non-synonymous SNPs being the most common DNA sequence variation associated with human diseases (32). Missense mutational effects can alter protein dynamics and cause human disease. Accurate evaluation of these effects can provide information on residue-specific roles in protein structure/function and dysfunction in the disease state. The current study advances the field of rare diseases, particularly Kleefstra syndrome, by implementing a computational biophysics approach and contrasting it with previous tools recommended by established guidelines.

The sequence-structure-function relationship has been established for all proteins, but molecular dynamics still need to be fully explored in genomic variation interpretation. To improve the assessment of genomic variations, we implemented a comprehensive computational approach incorporating multiple mechanism-based aspects of protein sequence, structure, and dynamics of EHMT1 for its mutational impact assessment. Structurally coordinated dynamics play an essential role in substrates binding and undergoing allosteric transitions while maintaining the native fold in catalytic enzymes (33, 34). Thus, dynamics-related protein-specific metrics can be reliable indicators of any protein function and dysfunction by disease mutations. Once identified, these metrics can be parameterized for each protein and domain. We analyzed each variant independently to show how curated missense variants may affect EHMT1 enzymatic activities. The selected metrics used in the current study served as effective measures of damaging impacts. The cross-correlation matrix of the individual scores was used to select more functionally relevant and effective metrics for each protein. Our findings affirm that the protein sequence-structure-dynamics-function relationships, molecular dynamic properties, and molecular structures have been conserved throughout evolution.

We hypothesize that universal structure-based metrics, such as folding/stability, global/local structural perturbation, and global/local energetic frustration, can be applied to all proteins and become part of the standard procedure for clinical functional impact analysis. On the other hand, MD-based metrics are not equally effective, and more congruent and functionally relevant MD-based metrics need to be identified for each protein. Moreover, a domain-wide analysis should be considered, especially when individual domains do not make any physical contact, because individual domains are independent folding units and functional modules.

Many proteins consist of several domains, and the same domain may appear in various proteins. Because members of the same domain family likely share the same evolutionary origin and perform similar molecular functions, the same set of effective MD-based metrics can be applied to the same domain family members. In many cases, long stretches of disordered regions connect individual domains and only individual domain structural information is available. Even when the multi-domain structures are available, individual domains can be isolated during data analysis and used for domain-specific analysis to gain more domain function-specific impact analysis. Subsequent collective analysis on multi-domain or functional oligomeric structures can provide additional mechanistic information, such as interdomain communication and cooperative functionality.

In conclusion, the current work provides important molecular-level insights into functional disruption by Kleefstra syndrome variants. Our data indicate that damaging variants of EHMT1 display mechanistic disruptions at either a structural or dynamics level or both, mainly concentrated around functional regions such as the active site and the substrate binding interface and the pre-SET region that contains the structural zinc ion cluster and the dimerization interface. On the other hand, tolerated variants are mostly found on the periphery or on the molecular surface, whose sequences are varied between EHMT1 and EHMT2. These findings should apply to not only EHMTs but also related SET domain-containing methyltransferases. Extended studies that use sequence paralogs and molecular dynamics will help validate our findings and improve the predictive value of these mechanistic-based comprehensive approaches. Furthermore, this comprehensive impact analysis should help annotate the pathogenicity of many different proteins that can be curated into the public archives of human genomic variations for clinical applications.

## MATERIALS AND METHODS

### Preparation of the initial structures

We constructed the two-domain structure containing both the SET and the ankyrin repeat domains by superposing the overlapping helix (982-998) of the individual domain structures (PDB access codes 3HNA and 6BY9) after considering the asymmetric contents of the crystal lattice. For the SET domain-only structure, we used the mono-methylated H3K9 peptide-bound form of the high-resolution (1.5Å) crystal structure (PDB access code 3HNA). The cofactor SAH was replaced with SAM to constitute the active form of the enzyme, and the seven missing residues in the flexible loop (965-971) were built using the Modeller program (35). The EHMT1-EHMT2 SET domain-only heterodimer was prepared by replacing its heterodimer partner from either homodimer structures (PDB access codes 3HNA or 5JJ0), which display a nearly identical dimerization binding mode. For missense variant analysis using these structures, substitutions were made within the Discovery Studio suite version 21.1 (Dassault Systèmes BIOVIA) by mutating the corresponding residue and selecting the side chain rotamer causing the least steric hindrance with the surrounding residues.

### Protein folding energy and stability calculation

We assessed the stability of the mutated protein by the variant-induced changes in folding energy (ΔΔG_fold_) using FoldX (36) and the Discovery Studio suite. We used the energy-minimized mutant structures for these calculations. In Discovery Studio, shifted amounts of protein stability (free-energy difference between folded and unfolded states) due to mutations were calculated at pH 7.4 using the energy-minimized wild-type structure and introducing each substitution for calculation. After the preparation phase, the initial structures of the wild type and the generated mutants were subjected to a two-stage minimization process before energy calculation. The predicted ΔΔGs, using both programs, are in good agreement (Supplementary Table S1 and Supplementary Fig. S4A-B).

### Global and local structure perturbation measurement

We measured the positional displacement of backbone atoms between the entire catalytic domains of wild-type and mutant (global) or only the atoms near the residue of interest between them (local). For local structure perturbation from the energy-minimized structures, any residues within a 10 Å radius of the mutation site were selected using PyMOL (Schrödinger, LLC) and calculated for least-squared RMSD of the backbone atoms between the wild type and the mutant using Coot (37). For global structure perturbation, entire backbone atoms were used for RMSD calculation between the structures.

### Frustration index calculation

The energy landscape of protein molecules can affect their biological behaviors. To evaluate how the energy is distributed in protein structures and how mutations or conformational changes shift the energy distributions, we measured the differences in energetic frustration in protein molecules using the Protein Frustratometer server (38). The *frustration index* measures how favorable a particular contact is relative to all possible contacts in that location. Sites of high local frustration often indicate biologically important regions such as binding or allosteric sites. The shift amounts were calculated by measuring the differences in the frustration indexes between the wild type and the mutant residues in both directions and summing up the differences. We tested the changes in either global (cumulative) or local frustration indexes as means of damaging impact scoring. We discovered that the global changes show better correlations with the sequence-based scores, which aligns with the multiple binding platforms used by this domain for various protein-protein interactions. Therefore, global changes were used for the overall impact scoring of the variants in the ankyrin repeat domain.

### Molecular Simulations

MD simulations were performed using the CHARMm36 all-atom-force-field (36) implemented in the Discovery Studio with a 2 fs time step. A simplified distance-dependent implicit solvent environment was used with a dielectric constant of 80 and a pH of 7.4. All MD simulations were conducted using periodic boundary conditions.

Models were energy minimized for 5,000 steps using the steepest descent followed by 5,000 steps of the conjugate gradient to relax the protein structure obtained under the stressed crystal environment. Each system of 10 replicates of wild type and each variant was independently heated to 300 K over 200 ps and equilibrated for 500 ps, followed by ten ns production simulation under NPT ensemble (100 ns total). Conformers were recorded every 10 ps to give 1000 frames for analyses per each mutation. This timescale is sufficient for side chain rearrangements in the protein’s native state and to facilitate local conformational changes. Total energy plots of the trajectories indicate that the systems can reach near equilibrium towards the end of the simulation. For final data analysis, one or two outliers (in some cases none) from each data set of 10 replicates that deviate from the rest in RMSD plots and might represent the minor and rarer form of conformations (altogether 14% of the entire data) were excluded from averaging, and only the last 500 frames that have reached the near minimum total energy state were used. From 10 ns MD simulation, trajectory files were analyzed for structural impact by root mean squared deviation (RMSD), root mean square fluctuation (RMSF), and other measures such as time-dependent molecular interactions, a radius of gyration (Rg), and solvent-accessible surface area (SASA). Trajectories were aligned to the initial WT conformation before analysis. RMSD and RMSF values were calculated at the residue level for all atoms using the tools available within Discovery Studio and the algorithms implemented in Microsoft Excel. Further analyses were conducted in the R programming language (39), leveraging the bio3d package (40). Molecular visualizations were generated using PyMOL.

### Time-dependent interaction energy calculation and distance monitoring

Molecular interaction-free energies were measured using Discovery Studio. This was done using the MD simulation trajectories and selecting the protein and the interaction groups of interest. Non-bonded interactions were monitored, and dynamic interaction energies (van der Waals and electrostatic energies) were calculated using the CHARMm36 force field and the implicit distance-dependent dielectric solvent model. Distance monitoring of the key catalytic components was also done within Discovery Studio by selecting those atoms of interest. These measurements were made for all 10 replicates and averaged for comparison with the wild type.

### Overall impact classification of the variant

We used a cross-correlation matrix among the scores as guidance to choose more effective and functionally relevant metrics for integration (Supplementary Text S1). For overall impact scoring, we Z-score scaled the individual scores onto a zero to one scale, commonly used by many sequence-based tools, and averaged them for the final scores (24). Finally, for variant classification, we used the suggested thresholds for sequence-based prediction tools for overall impact scoring as guidelines and re-classified the variants based on the meta-scores (0-0.3: tolerated, 0.3-0.4: uncertain, and 0.4-1.0: damaging). This results in a balanced number of variants in each category. As a result, known damaging variants and gnomAD healthy population variants belong to the respectively expected category. Likewise, the proper threshold values were chosen for molecular fitness scores (without the sequence-based scores) based on the suggested values for sequence-based prediction tools. Any variants predicted to be damaging at either molecular aspect (structure or dynamics), yet the overall meta-scores below 0.4 have been assigned as VUS. Similarly, any variants that fall below the threshold value at either aspect, yet collectively the overall meta-scores exceed 0.4 have been assigned as damaging (but not further classified as either a structural or dynamic variant).

## Supporting information

Combined supporting information

## List of abbreviations

COSMIC: Catalogue of Somatic Mutations in Cancer
dbSNP: Single nucleotide polymorphism database
DV: Dynamic variant
EHMT1: Euchromatic histone lysine methyltransferase 1
GLP: G9a-like protein
gnomAD: genome aggregation database
KDM6A: Lysine-specific demethylase 6A
MD: Molecular dynamics
OMIM: Online Mendelian inheritance in man
PDB: Protein data bank
Rg: Radius of gyration
RMSD: Root mean square deviation
RMSF: Root mean square fluctuation
SAH: S-adenosylhomocysteine
SAM: S-adenosylmethionine
SASA: Solvent-accessible surface area
SDV: Structural and dynamic variants
SET: Su(var)3-9, Enhancer-of-zeste and Trithorax
SNP: Single nucleotide polymorphism
SV: Structural variant
VUS: Variant of uncertain (unknown) significance

## Data availability

MD Simulation data is available upon request.

## Acknowledgments and funding information

This work was funded by the Advancing a Healthier Wisconsin Endowment and the 501c3 charitable organization Harmony 4 Hope to the Precision Medicine Simulation Unit of the Mellowes Center for Genomic Sciences and Precision Medicine at the Medical College of Wisconsin (to RU). This work was also supported in part by NIH grants R35GM128840 (to BCS) and R01DK052913 (to RU and GL).

## Authors’ contributions

RU devised the project, the main conceptual ideas, and the proof outline. RU, MTZ, SDJ, and YC designed the computational framework and analyzed the data. DRJ, BCS, BFV, AJM, and GL discussed the results, provided critical feedback, and helped shape the research and analysis. YC, MTZ, and RU were major contributors to writing the manuscript. All authors read and approved the final manuscript.

