## Supplementary material for "A Multi-Layered Computational Structural Genomics Approach Enhances Domain-Specific Interpretation of Kleefstra Syndrome Variants in EHMT1": Combined supporting information

Young-In Chi, Salomão D. Jorge, Thiago M. De Assuncao, Elise N. Leverence, Brian C. Smith, Brian F. Volkman, Angela J. Mathison, Gwen Lomberk, Michael T. Zimmermann, and Raul Urrutia

#### Supplemental movie

**Movie M1.** EHMT1 wild type SET domain MD simulation (10 ns) animation. The bound H3 peptide and the SAM cofactor are shown as ball and stick models, while the protein is shown as ribbons.

#### Supplemental tables

**Table S1.** Spreadsheet of current annotations and reclassification based on individual scores, scaled Z-scores, and integrated overall impact scores. The overall scores are made by summing up the scaled Z-scores from each metric with equal weight and averaged. The damaging variants are further classified as structural variants (SV), dynamic variants (DV), or structural & dynamic variants (SDV) based on their 'molecular-fitness' scores.

#### Supplemental text

#### Text S1. R script used for generating the cross-correlation matrix

```
library(readxl)
library(corrr)

df <- read_excel("impact-score.xlsx")
df

df1 <- select(df, SNPsGo, MutPred2, PROVEAN, MutationAssessor,
PolyPhen2, Rhapsody, Missense3D, FoldX, DynaMut, iStable,
StabilityEnergy, GlobalPerturb., LocalPerturb., pKashift,
LocalpKashift, TriadInteraction, H3ZnInteraction, RMSD.median.diff,
Inv.RMSF.Scorr, Rg, SASA)

df1[df1==""]  <- NA
str(df1) # to review the data structure

df2 <- select(na.omit(df1), SNPsGo, MutPred2, PROVEAN,
MutationAssessor, PolyPhen2, Rhapsody, Missense3D, FoldX, DynaMut,
iStable, StabilityEnergy, GlobalPerturb., LocalPerturb., pKashift,
LocalpKashift, TriadInteraction, H3ZnInteraction, RMSD.median.diff,
Inv.RMSF.Scorr, Rg, SASA)
str(df2)

library(corrplot)

spearman_corr <- cor(df1, method = "spearman")

corrplot(spearman_corr, type = "upper",
         tl.col = "black", tl.cex = 0.75, font = 2, method = "square",
addCoef.col = "black",
         number.cex = 0.5,
         diag = FALSE)
```

**Figure S1**

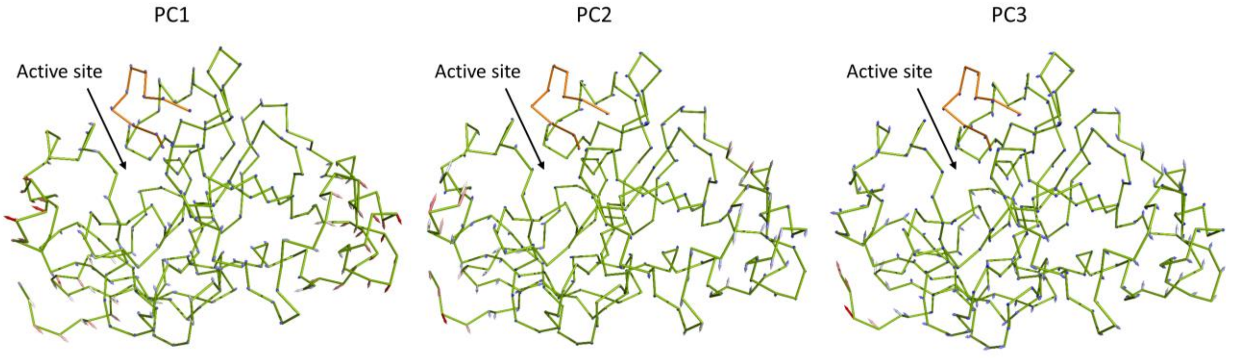


**Figure S1.** Porcupine plots of the trajectory representing the essential dynamic motions in each principal component during MD simulation of the wild type EHMT1 SET domain. The length of the arrows represents the motion magnitude and the pointing of the arrow indicates the direction. The catalytic SET domain movements display high mobility in the pre-SET region containing a dimerization interface, and relatively rigid motions in the active site and substrate binding site.

**Figure S2**

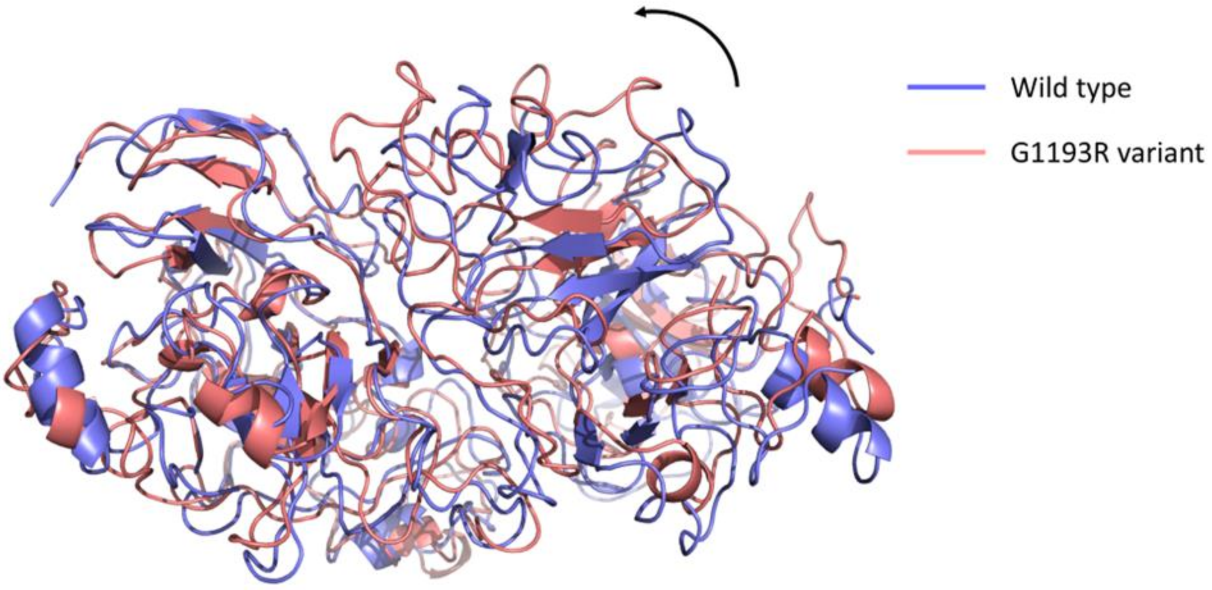


**Figure S2.** Alterations in overall RMSD due to a rotation with respect to each monomer (G1193R variant as an example). The relative orientation between the monomers is expected to play a critical functional role.

#### Figure S3

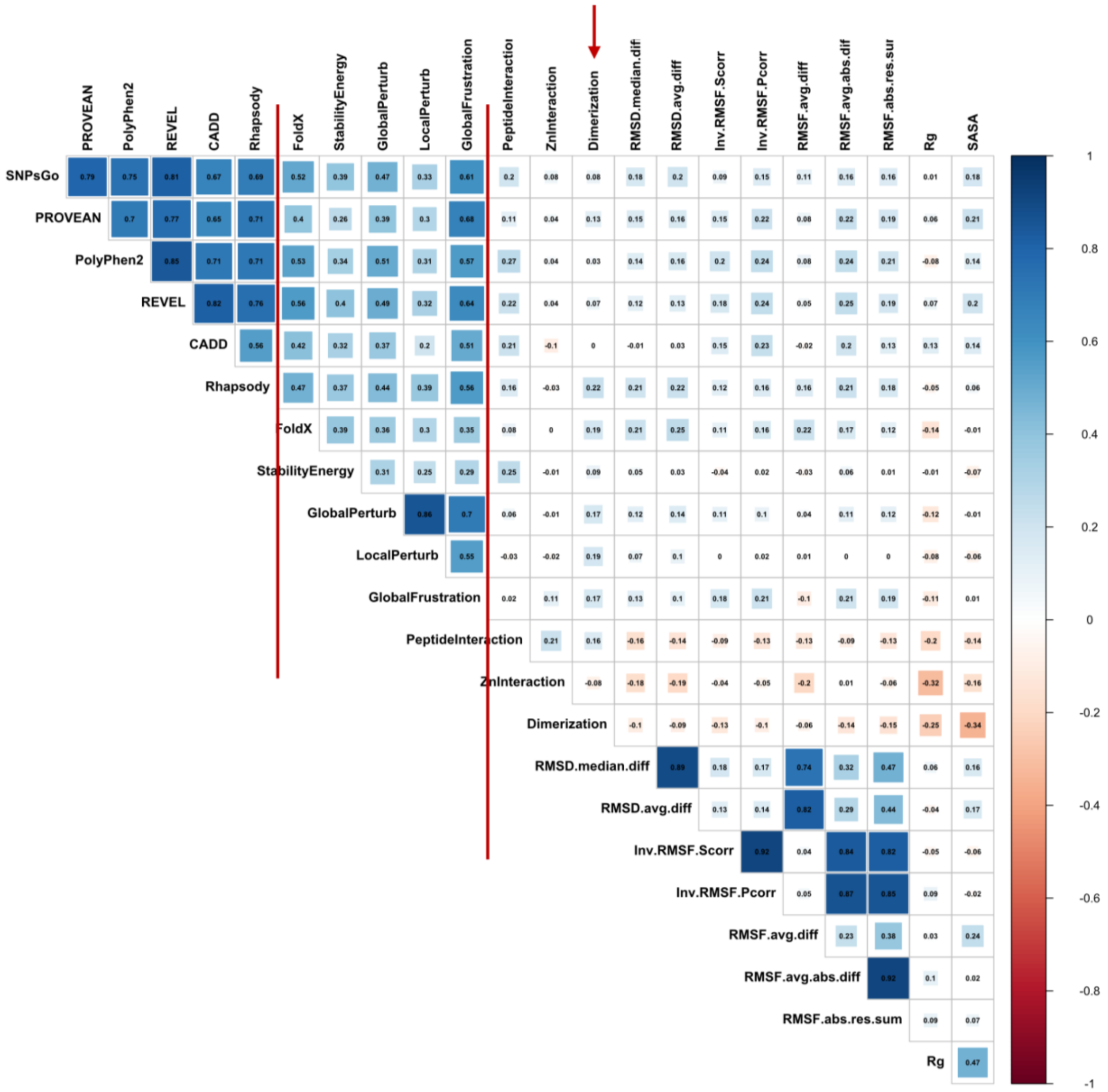


**Figure S3.** Cross-correlation matrix with various measures of universal and protein-specific metrics from the EHMT1:EHMT2 SET-domain heterodimer. Among the protein-specific MD-based scores, H3K9me peptide interaction energies show noticeable correlations as expected while dimerization interaction energies (indicated by an arrow) show weaker correlations with the sequence-based scores.

### Figure S4

A

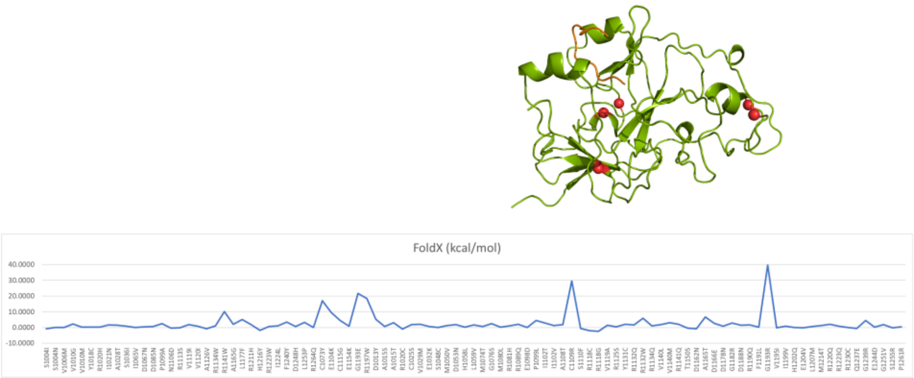


B

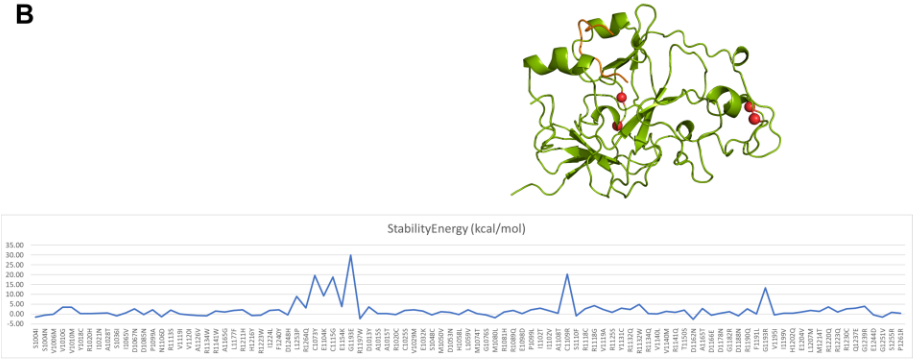


C

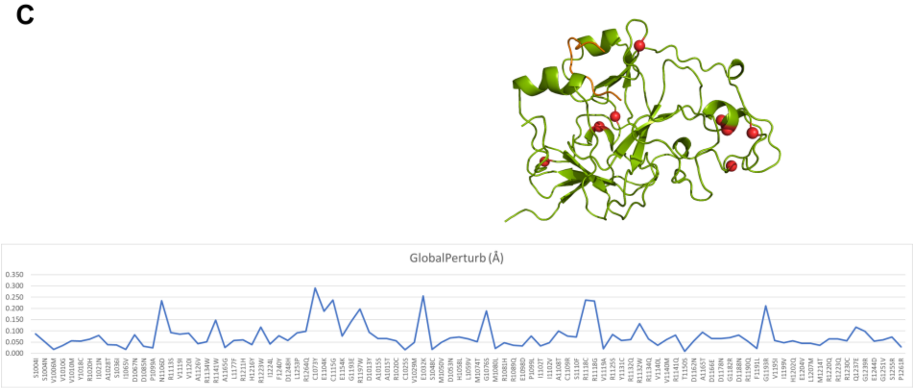


D

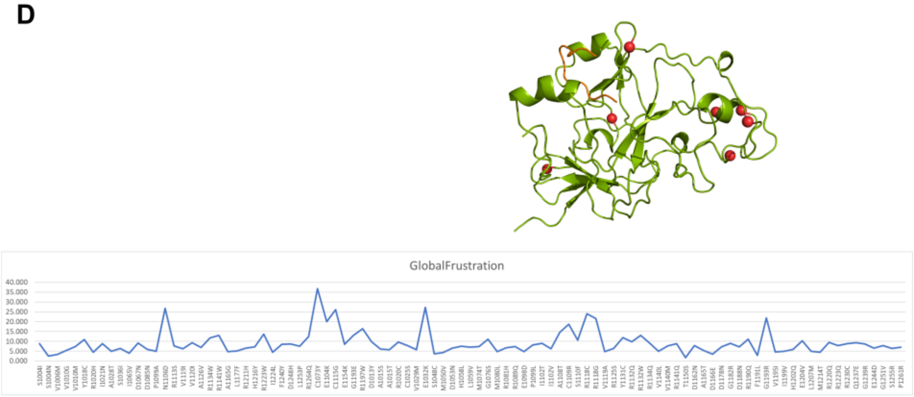

### Figure S4 (cont.)

E

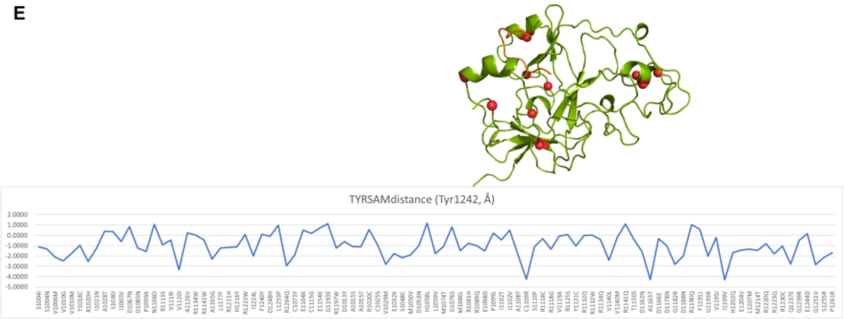


F

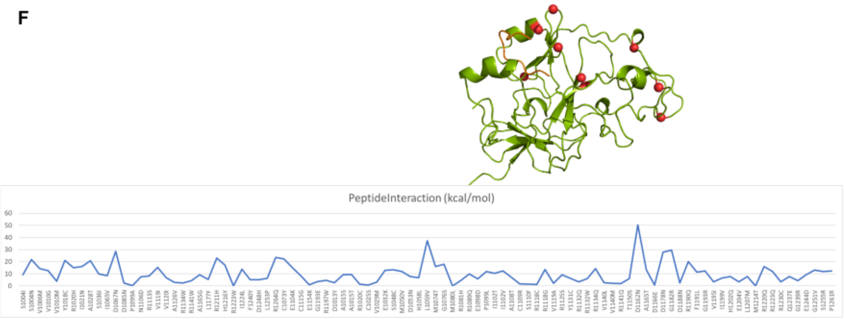


G

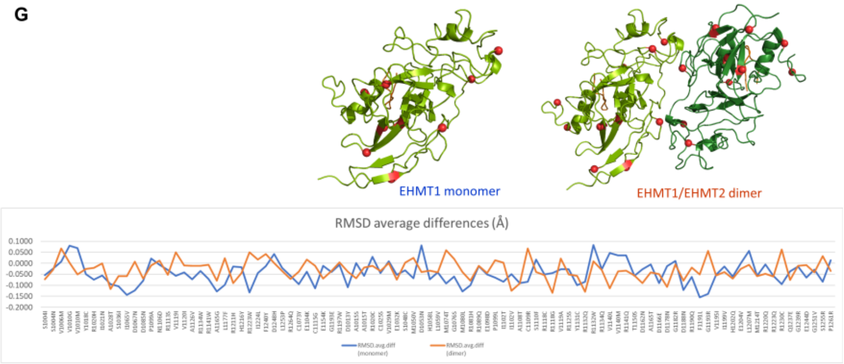


H

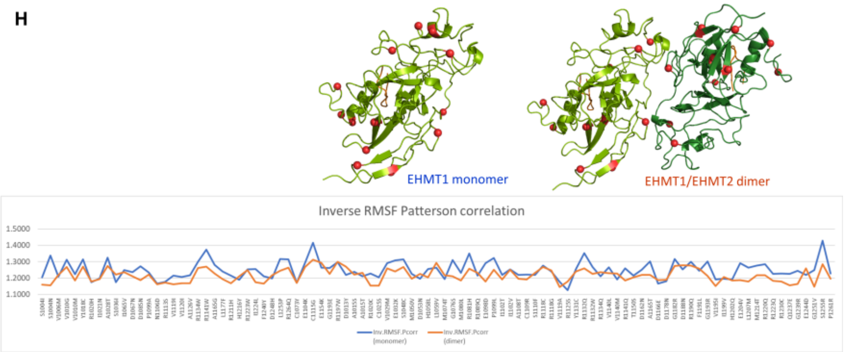


I

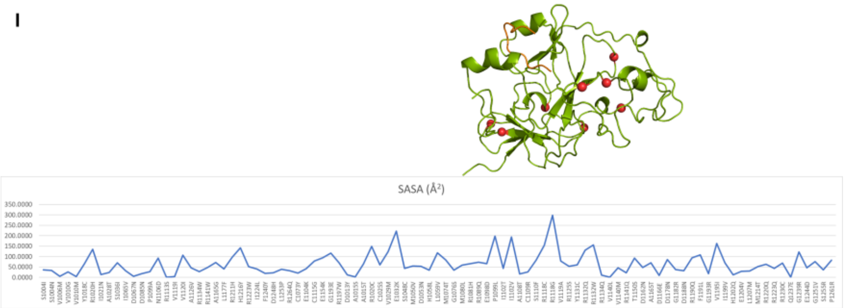

**Figure S4.** Plots of individual damaging scores for each metric and mapping the most damaging variants onto the structure. The variants on the horizontal axis are in the same order as the ones in Supplementary Table S1. Structure-based metrics (A-D) and MD-based metrics (E-I) are shown. For RMSD and RMSF, both monomeric and heterodimeric results are shown for comparison.
